# Arginine multivalency stabilizes protein/RNA condensates

**DOI:** 10.1101/2021.04.22.440959

**Authors:** Matteo Paloni, Giovanni Bussi, Alessandro Barducci

## Abstract

Biomolecular condensates assembled through liquid-liquid phase separation (LLPS) of proteins and RNAs are currently recognized to play an important role in cellular organization. Their assembly depends on the formation of a network of transient, multivalent interactions between flexible scaffold biomolecules. Understanding how protein and RNA sequences determine these interactions and ultimately regulate the phase separation is an open key challenge. Recent *in vitro* studies have revealed that arginine and lysine residues, which are enriched in most cellular condensates, have markedly distinct propensities to drive the LLPS of protein/RNA mixtures. Here, we employ explicit-solvent atomistic Molecular Dynamics (MD) simulations to shed light on the microscopic origin of this difference by investigating mixtures of polyU oligonucleotides with either polyR/polyK peptides. In agreement with experiments, our simulations indicate that arginine has a higher affinity for polyU than lysine both in highly diluted conditions and in concentrated solutions with a biomolecular density comparable to cellular condensate. The analysis of intermolecular contacts suggests that this differential behavior is due to the propensity of arginine side chains to simultaneously form a higher number of specific interactions with oligonucleotides, including hydrogen bonds and stacking interactions. Our results provide a molecular description of how the multivalency of the guanidinium group enables the coordination of multiple RNA groups by a single arginine residue, thus ultimately stabilizing protein/RNA condensates.

## Introduction

Biological condensates that are not bound by membranes have recently emerged as central elements in the organization and regulation of cellular functions^1–4^. An increasing amount of evidence is showing that these membraneless organelles (MLOs) are formed by liquid-liquid phase separation (LLPS) of complex mixtures of proteins and nucleic acids^5–7^. In this process, intrinsically disordered proteins/regions (IDP/IDRs) play a key role since their structural flexibility allows the formation of a network of multivalent, transient interactions that stabilize the condensed phase. In several cases, a single IDP/IDR component has been shown to undergo liquid-liquid demixing *in vitro* and to form assemblies similar to cellular condensates, thus providing tractable model systems to investigate the physico-chemical principles governing the LLPS^8,9^. In the last few years, the combination of experimental and theoretical investigations has provided some insight on how the patterning of charged and aromatic residues in IDP/IDR sequences modulates the intra- and inter-molecular interactions and ultimately determines the phase behavior^8,10^. Conversely, the molecular grammar governing the assembly of protein/RNA condensates has been less studied, even if RNAs are abundant in most cellular MLOs and they modulate their assembly, physical properties and biological function^11^. In the cell, this complex interplay of protein and RNA components arises from a large variety of homotypic and heterotypic interactions, which often include sequence-dependent recognition of RNA molecules by RNA-binding modules in the protein components and/or specific RNA/RNA interactions. On more general grounds, the negative charge of RNA is the most basic feature that can promote protein-RNA interactions and drive condensate assembly through electrostatically-driven complex coacervation. Remarkably, even simple model systems, such as RNA homopolymers and low-complexity polypeptides, have been shown to give rise *in vitro* to complex coacervates with multilayered architectures and a broad range of material properties in a sequence-dependent manner^12^.

While we are still far from fully understanding how nucleotide and amino-acid sequences determine the assembly and properties of condensates, recent evidence indicates that lysine and arginine residues have markedly different propensities to drive the LLPS of IDPs^10,13,14^ and IDP/RNA mixtures^15–17^. This finding is particularly remarkable considering that both these basic amino acids are positively charged at physiological pH and are significantly enriched in the low-complexity protein domains that drive the assembly of MLOs. Notably, arginine-rich peptides have been shown to phase separate with polyU RNA at lower concentration than lysine-rich peptides and to form coacervates that are less dynamical^15^. This differential behaviour is more pronounced if one compares the demixing propensities of polyK/polyU and polyR/polyU mixtures and the viscosities of the resulting condensates^16^. Furthermore, NMR spectra revealed that lysine/RNA-interactions differ from arginine/RNA-interactions resulting in distinct molecular environments in protein/RNA droplets^15^. Nevertheless, a comprehensive microscopic picture is still lacking and multiple molecular mechanisms have been invoked to justify these results, including differences in protonation equilibria, H-bonding network, hydrophobicity and/or capability of establishing pi-pi and cation-pi interactions with nucleobases^12,15^. In this work, we take advantage of Molecular Dynamics (MD) simulations to investigate the interactions of arginine and lysine side chains with RNA molecules in biomolecular condensed phases. MD simulations are a promising tool for characterizing the molecular determinants of LLPS and the structural properties of biomolecules within condensates, circumventing the limitations of most experimental approaches in structural biology. In particular, Coarse-Grained (CG) simulations at various levels of resolution have been recently used for studying the assembly and properties of RNA/protein condensates^18–20^. While CG models are necessary to tackle the time and length scales associated with LLPS processes, these models are based on phenomenological energy functions and therefore cannot fully capture the microscopic driving forces underlying phase separation. Therefore, here we adopted explicit-solvent atomistic MD to exhaustively characterize the interactions of polyR/polyK peptides with polyU oligonucleotides at various concentrations, building on our recent work on the molecular interactions underlying the phase separation of DDX4 protein^21^.

## Results

We investigated the behaviour of polyR/polyU and polyK/polyU mixtures by using as molecular models five-residue long arginine and lysine peptides (R_5_ and K_5_, respectively) together with five-residue long polyuridylic acid RNA fragments (U_5_). This choice is justified by several factors: *i)* the good accuracy of last-generation force fields in reproducing the structural ensemble of small oligonucleotides^22^, *ii)* the potential sampling difficulties associated to slower conformational dynamics of long peptides/oligonucleotides^23^ and, most importantly, *iii)* the experimental observation that the differential behaviour of polyR/polyU and polyK/polyU is not dependent on molecular size^16^.

We first characterized the relative affinity of R_5_ and K_5_ with U_5_ in highly-diluted conditions by performing unbiased MD simulations of a single peptide chain together with one oligonucleotide chain. MD trajectories (see Methods) revealed multiple binding/unbinding events although kinetic bottlenecks hinder an accurate determination of the thermodynamic stability of the peptide/nucleotide complexes, especially in the case of R_5_/U_5_ system, on the microsecond time scale (Fig.1A,B). To overcome these sampling difficulties, we relied on REST2 Hamiltonian replica-exchange method (see Methods) that has been shown to greatly accelerate the conformational sampling for solvated biomolecules^24^. Adopting this approach, we could reliably evaluate the stability of the peptide/oligonucleotide complexes by determining the free energy profile (FEP) as a function of the intermolecular interaction surface for both R_5_/U_5_ and K_5_/U_5_ (Fig. 1C). Both profiles display a free-energy minimum corresponding to unbound states (interaction surface = 0) and a large basin associated to a variety of structurally-heterogeneous complexes, as expected for a dynamical, electrostatically-driven interaction between two flexible biomolecules. Comparison of the FEPs indicate that in highly-diluted conditions R_5_ binds to U_5_ more strongly than K_5_ and it forms complexes characterized by a larger interaction surface, as it was already hinted at by the unbiased simulations. A more quantitative analysis of REST simulations provided a rough estimate of the complex dissociation constants, which differ by more than one order of magnitude (K_d_=23 μM for R_5_/U_5_, and K_d_=850 μM for K_5_/U_5_, see Methods).

**Fig. 1-.**
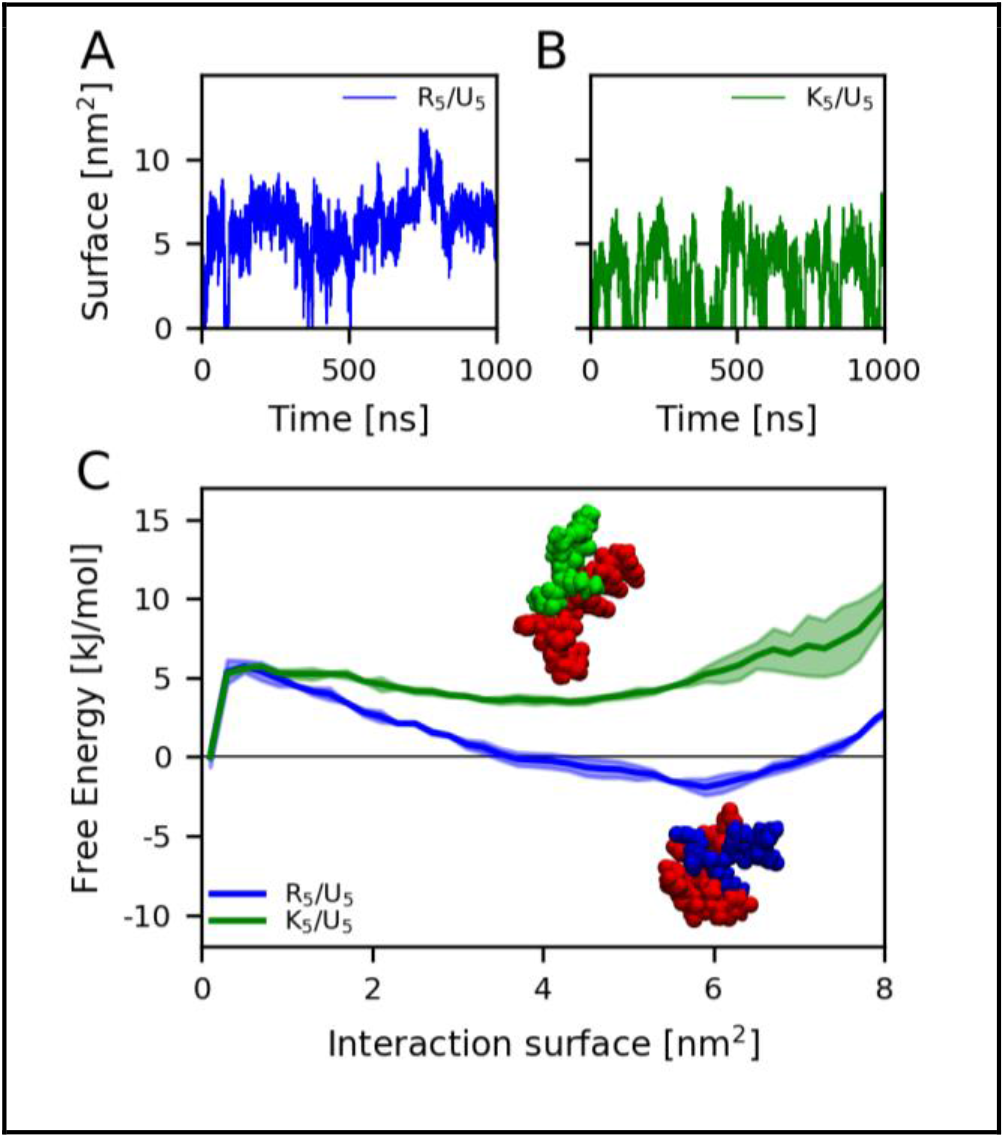
Time series of the interaction surface from unbiased simulations of polyR/polyU (A) and polyK/polyU (B) systems. (C) Free energy profile of polyR/polyU and polyK/polyU interactions as a function of the interaction surface between the two molecules. Shaded areas correspond to the standard deviation between the first and the second half of the simulation. Insets are representative configurations of the systems at the local minimum of the free-energy profile corresponding to the bound state.

We then sought to characterize the molecular interactions in conditions mimicking the dense protein/RNA coacervates. Unfortunately, the actual molecular composition of polyK/polyU or polyR/polyU condensed phases, in terms of RNA/protein ratio and water content, is yet to be determined experimentally. Phase coexistence simulations^25,26^ might in principle provide access to this information but their applications at the atomistic level would require a prohibitive computational effort even in the case of small peptides/oligonucleotides. To circumvent this difficulty, we decided to focus here on R_5_/U_5_ and K_5_/U_5_ mixtures with a 1:1 molecular ratio and to characterize their behavior in a range of concentrations comparable to those determined for *in vitro* model biomolecular condensates. For this reason, we simulated peptide/oligonucleotide mixtures with biomolecular densities of approximately 125, 250, and 500 mg/ml, which correspond to diverse volume packing fractions (Fig. 2A, B, and C, and table S1). Even in these highly crowded conditions, peptides and nucleotides form dynamical intermolecular contacts, without formation of irreversible complexes (Fig. S1). We first inspected the overall arrangement of the biomolecules in the various systems by determining the Radial Distribution Function (RDF) of the intermolecular distances between all the biomolecular heavy atoms (Fig. 2D, E, and F). The comparison of these intermolecular RDFs with the distributions corresponding to random arrangements revealed noticeable attractions between the biomolecules, independently of the concentration and the nature of the peptides. These favourable interactions result in an increased average local density in the nanometer scale that is particularly evident at lower concentrations and more pronounced for R_5_/U_5_ systems, suggesting stronger intermolecular binding. In order to dissect the molecular driving forces, we then analysed the average number of protein-protein, protein-RNA and RNA-RNA contacts per molecule (see Methods and Fig. 1G, H, and I). In all the systems, heterotypic interactions between peptides and oligonucleotides represent the largest contribution to intermolecular contacts, as expected due to the electrostatic attraction between the oppositely-charged moieties. Consequently, homotypic interactions play a more limited role. In particular, protein-protein contacts are extremely limited in all the mixtures, due to the electrostatic repulsions between the positively-charged peptides. This effect is less pronounced in the case of RNA-RNA interactions, which account for a sizable fraction of the intermolecular contacts, possibly due to the smaller charge density of U_5_ oligonucleotides that are larger and have a smaller net charge than R_5_/K_5_ peptides. Nevertheless, the electrostatic interactions cannot account for the higher interaction propensity of polyR with respect to polyK. Homotypic polyR-polyR contacts are indeed more frequent than polyK-polyK ones at all concentrations, as expected due to the well-recognized tendency of arginine side chains to form favorable stacking interactions. Even more strikingly, R_5_ peptides display a significantly higher propensity to bind oligonucleotides U_5_ at all the concentrations, even if the difference between the number of heterotypic contacts is mitigated in denser, more crowded systems (∼50% relative increase at 125 mg/ml with respect to ∼25% to 500 mg/ml). Next, we evaluated the mobility of peptide and oligonucleotide chains in the studied systems by estimating the diffusion coefficients for the various species at different concentrations (Fig. 2J, K, and I). The results indicate that molecular diffusion is strongly dependent on the concentration as diffusion coefficients decrease roughly by two orders of magnitude between mixtures at 125 mg/ml and those at 500 mg/ml. Beyond this overall trend, the analysis suggests that the oligonucleotides are slightly less mobile than polypeptides, consistently with their larger size and molecular weight. The stronger intermolecular interactions and higher local densities observed in R_5_/U_5_ systems slow the diffusion of all their components by a factor of ∼2 with respect to K_5_/U_5_ at equivalent concentration.

**Fig.2.**
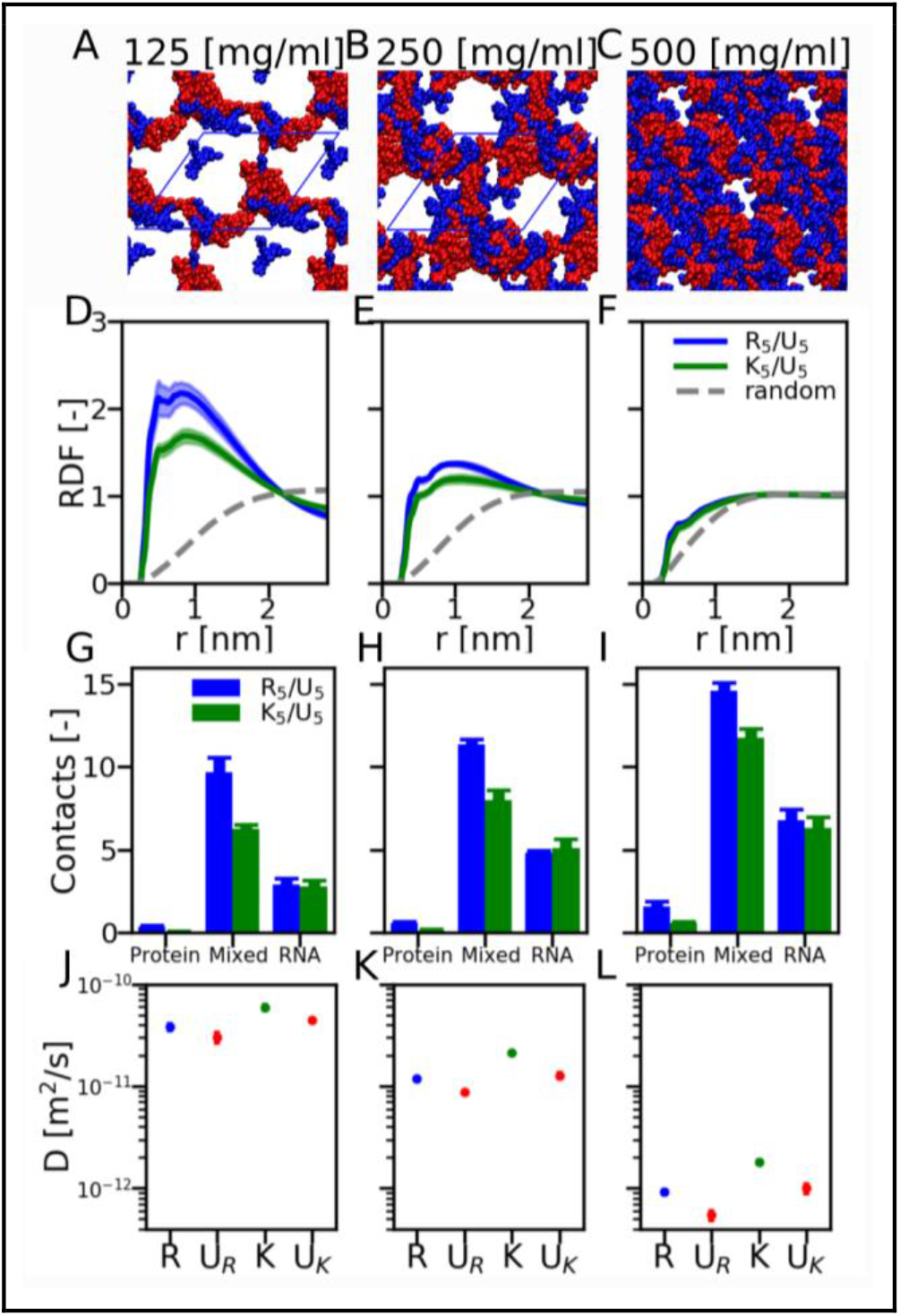
(A,B,C) Representative snapshots from the MD simulations of poly-R/poly-U simulations at the three concentrations investigated: 125 mg/ml (A), 250 mg/ml (B), and 500 mg/ml (C). The periodic box is shown in blue. (D,E,F) Radial distribution function (RDF) of the intermolecular heavy-atom pair distances between all protein and RNA atoms. Dashed lines are the RDFs for random configurations, and exhibit a depletion at short distances due to the excluded volumes of the two molecules. (G,H,I) Intermolecular contacts per chain for Protein-Protein, Protein-RNA, and RNA-RNA interactions. (J,K,L) Diffusion coefficients of peptide (R and K) and oligonucleotide (U_R_ and U_K_, corresponding to the poly-R and poly-K simulations, respectively) molecules.

In order to get a deeper insight on the microscopic determinants of protein-RNA interactions and to elucidate the different behaviour between arginine and lysine residues, we first broke up the contributions of backbone and side chain groups to the number of protein-RNA contacts (Fig. 3A,B). This analysis indicated that backbone-mediated heterotypic interactions are similar for R_5_ and K_5_ and that the different interaction propensity with oligonucleotides mostly originates from contacts with the structurally-diverse side chains. Thus, we focused on the latter and we determined the RDF between the peptide side chains and the various chemical groups in the oligonucleotides, *i*.*e*. phosphate, ribose, and base (see Methods). Overall, RDFs corresponding to R_5_/U_5_ and K_5_/U_5_ systems exhibit significant differences both in the position and the intensity of the peaks, immediately suggesting distinct binding modes for the arginine and lysine side chains. In both systems, the direct interaction with the phosphate group is characterized by a major peak at <0.5 nm (Fig. 3C,D upper panel), which corresponds to the formation of Hydrogen bonds (H-bonds) between phosphate oxygens and the basic groups of the side chains (Fig. 3C,D insets and lower panel). Remarkably, the average number of simultaneous H-bonds formed by arginine is approximately twice as much as those formed by lysine. The interpretation of the side chain interactions with the ribose (Fig. 3 E,F upper panel) is less straightforward. The broad peak between 0.4 and 0.7 nm observed for both polyK and polyR indicate a variety of binding modes characterized by similar distances, which can be ascribed to the complex shape of the sugar group. For both side chains, a fraction of these interactions implies the formation of H-bonds with the hydroxyl group of the sugar (Fig. 3E,F insets and lower panel). Conversely, RDFs suggest that arginine and lysine interactions with RNA bases differ significantly (Fig. 3G,H upper panel): while lysine presents a single peak at a distance of ∼0.6 nm, corresponding to H-bonding with the carbonyl oxygens of the nucleobase ring, arginine side chains also exhibit an alternative binding mode at lower distances, representing a stacking arrangement of the nucleobase and the planar guanidinium group (Fig. 3G,H insets and lower panel) that is reminiscent of the stacking interactions between arginine and aromatic amino acids.

**Fig.3.**
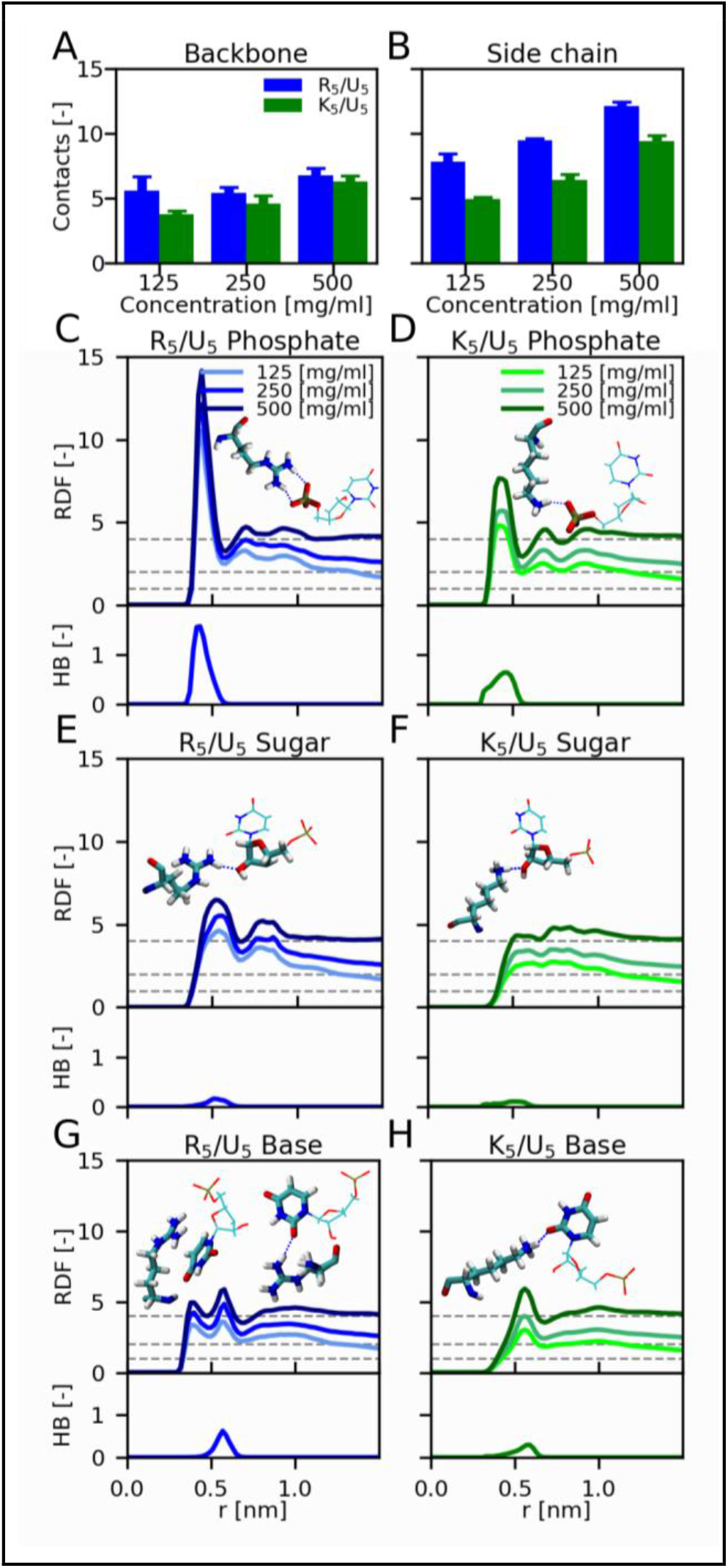
(A,B) Number of contacts per chain formed by backbone (A) and side chain (B) atoms of Arg and Lys with RNA at different concentrations. (C,D,E,F,G, and H) Radial distribution function (RDF) between the CZ/CE atoms of R_5_/K_5_ and the centers of geometry of the RNA groups. RDFs are normalized by the ratio of the concentration of the system at a reference value (125 mg/ml) (upper panels). Dashed horizontal lines show the expected value at large distance for the simulated concentration. Number of hydrogen bonds (HB) formed by Arg and Lys side chains as a function of the distance between CZ/CE atoms and centers of geometry of RNA groups (lower panels).

The previous analysis highlighted that both arginine and lysine side chains can form a variety of H-bonds with the diverse chemical groups of oligonucleotides. In order to rationalize the different behaviour of polyR and polyK, we thus calculated the average number of H-bonds formed simultaneously by a single amino acid, limiting the average to residues that are in direct contact with polyU to reduce its dependence on the system density (Fig. 4A). This analysis reveals that each arginine forms approximately twice as many H-bonds as lysine side chains. This result does not depend on the overall system concentration and it can be attributed to the structural differences between the amino-acid side chains and to the availability of more hydrogen donor atoms in arginine side chains. Furthermore, the guanidinium groups can contribute to the intermolecular contact network also through stacking interactions with the nucleotide bases as suggested by RDF. The average number of these interactions formed by each arginine is significantly smaller than the number of H-bonds but far from being negligible (Fig. 4B). Therefore, the two-pronged and planar arginine side chain enables the simultaneous formation of a larger and more diversified set of protein-RNA interactions than the lysine side chain. This multivalency is pictorially exemplified by some representative conformations extracted from MD trajectories where guanidinium groups coordinate multiple bases via H-bonds and stacking interactions (Fig. 4C, left panel) or bridge. In fact, various examples can be identified of guanidinium groups coordinating multiple bases via hydrogen bonds and pi-pi interactions (left panel of Fig. 4C), or forming bridge interactions between phosphate and base groups (central and right panel of Fig. 4C).

**Fig.4.**
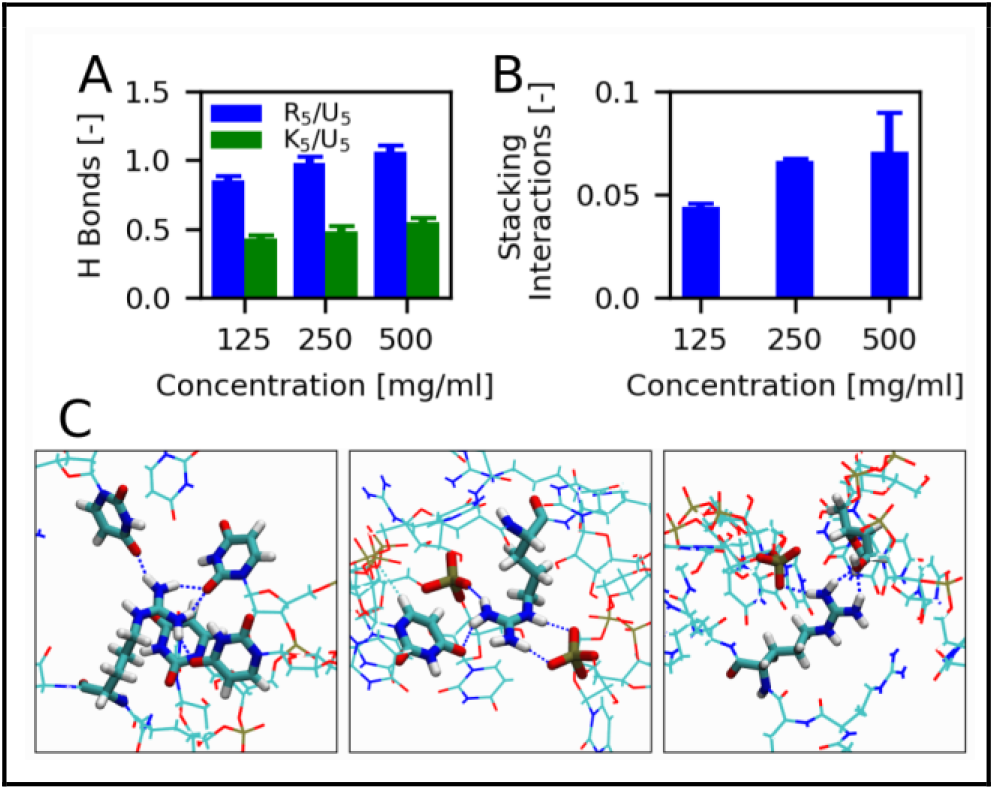
(A) Number of hydrogen bonds per amino-acid side chain in contact with RNA. (B) Number of stacking interactions per arginine side chain in contact with RNA. (C) Representative configurations extracted from MD simulations of R_5_/U_5_ showing the coordination of multiple RNA groups by arginine side chains.

## Discussion

In this work we used explicit-solvent atomistic simulations to shed light on the molecular determinants of protein-RNA LLPS by investigating the distinct role of arginine and lysine in determining the demixing propensity and the condensate properties. To this aim, we first characterized the interactions of R_5_/K_5_ peptides with U_5_ oligonucleotides in highly-diluted conditions and observed that both peptides form dynamical fuzzy complexes, whose thermodynamic stabilities were accurately determined by means of an enhanced sampling method. Our calculations indicated that R_5_ has a higher affinity than K_5_ for U_5_, with a difference of more than one order of magnitude in the relative dissociation constants, in good agreement with the binding affinities determined for longer polyU and polyK/polyR systems by fluorescence correlation spectroscopy experiments^16^. Reassured by this first result, we then aimed at elucidating the interactions between peptides and oligonucleotides in the dense protein/RNA condensates. To achieve this goal, we took inspiration from our recent results^21^, suggesting that atomistic MD simulations of peptides at high concentration can recapitulate fundamental aspects of protein LLPS. Therefore, we performed multiple MD simulations of highly concentrated R_5_/U_5_ and K_5_/U_5_ mixtures in a range of biomolecular concentrations (125-500 mg/ml) that is comparable to those estimated *in vitro* for biomolecular condensates^8,9,27^

Even if this approach cannot provide an exhaustive picture of the condensate assembly and composition, it partially circumvents the difficulties of performing phase coexistence simulations with computationally-demanding atomistic models and it can unravel the molecular interactions underlying the LLPS process. We observed that all the simulated peptide/nucleotide mixtures display strong non-ideal behaviour due to overall attractions between the biomolecules, resulting in significant local density fluctuations in the nanometer scale. This effect was more pronounced in polyR/polyU mixtures consistently with the higher phase-separation propensity experimentally determined for this system^16^ and with mutagenesis experiments on protein/RNA condensates^15,17^. The inspection of intermolecular contacts revealed that polyR peptides always form more homotypic interactions than polyK, reflecting the propensity of arginine side chains to self-interact through the stacking of the guanidinium groups, as already observed in crystal structures^28^ and atomistic simulations^21^. PolyR peptides also displayed a significantly higher propensity to bind oligonucleotides, further indicating that, other than the net charge, the structural differences between arginine and lysine amino acids affects their ability to interact with RNA molecules. Molecular diffusion was found to be strongly dependent on the system concentration with a decrease of almost two orders of magnitude between the diffusion coefficients of oligonucleotides and peptides at 125 mg/ml and those at 500 mg/ml. Furthermore, the stronger interactions in polyR/polyU mixtures resulted in a factor of ∼2 between the diffusivity of arginine and lysine peptides at the same concentration. This analysis suggests that the impressive difference in the viscosity of polyR/polyU and polyK/polyU condensates recently measured by microrheology experiments^16^ could be justified by a limited variation in their density.

MD trajectories indicate that both arginine and lysine side chains form a variety of distinct intermolecular contacts with polyU, which encompass H-bonding with the oxygens of the negatively-charged phosphates, the hydroxyl oxygen of the ribose and the carbonyl oxygens of the base, as well as guanidinium-uracil stacking in the case of arginine. All these dynamical interactions in peptides/nucleotides mixtures closely resemble those observed in X-ray structures of stable protein/RNA complexes^29,30^.

A defining aspect of LLPS is the presence of multivalent, transient interactions between biomolecules that stabilize the condensed phase^2,6^. In this respect, while both polyK and polyR can establish H-bonding networks with polyU, we found that arginine side chains are on average involved in twice as much H-bonds as those formed by lysine, thanks to the availability of more hydrogen donor atoms and a favorable geometric complementarity of guanidinium and phosphate groups^15^. Arginine multivalency is further reinforced by its capability of forming additional stacking interactions with nucleobases, although their frequency was rather limited in our simulations. Although we cannot exclude an underestimation of π-π and cation-π interactions by our classical force field, a recent structural analysis^31^ of protein/RNA complexes suggests that arginine and uracil have a limited propensity to form stacking interactions, in agreement with our findings. Altogether, the versatility of arginine guanidinium groups may eventually lead to complex binding patterns where a single arginine side chain coordinates several RNA groups, and possibly multiple oligonucleotide chains. These multivalent interactions are reminiscent of arginine forks, which are a widespread structural motif in protein/RNA complexes according to the extensive analysis of protein structural databases^29^. In conclusion, our results suggest that the peculiar structural features of arginine side chain play a key role in the formation of multivalent interactions with model unstructured RNAs, such as polyU oligonucleotides, thus rationalizing the important role of this amino acid in protein/RNA phase separation. Nevertheless, the observation that protein/RNA coacervates formed with various homonucleotides have distinct rheological properties^12^ strongly advocates for future investigation aimed at elucidating the role of RNA composition and sequence in protein/RNA condensation. In this respect, we are confident that atomistic MD simulations, possibly supplemented by enhanced sampling techniques, have the potential to substantially contribute to the definition of a comprehensive molecular grammar of protein/RNA phase behavior, extending pioneering investigations focusing on mRNA-protein complementarity^32^.

## Materials and methods

### Unbiased MD simulations

R_5_ and K_5_ peptides were generated in extended structure with the LEaP program in AmberTools16^33^, while U_5_ oligonucleotides were built in A-form using Chimera 1.14^34^. Peptide molecules were capped with ACE and NME groups to avoid artificial effects induced by the charged termini. Initial configurations for all simulations were generated by inserting peptide molecules and RNA fragments in random positions and orientations using the gmx insert-molecules tool in Gromacs 2018.3^35^. Simulations of systems at 125, 250, and 500 mg/ml were performed in rhombic dodecahedron boxes of ∼145 nm^3^, with respectively 10, 20, and 40 solute molecules equally divided in peptide and RNA chains, while a smaller box of ∼88 nm^3^ was used for the simulations of systems in highly-diluted conditions with only one molecule per species (∼40 mg/ml). However, in the latter systems, the presence of only two molecules within a large simulation box (side ∼5nm) naturally prevents homotypic as well as heterotypic interactions involving more than one polypeptide and one oligonucleotide, thus mimicking infinite dilution. Gromacs 2018.3^35^ was used to run all unbiased MD simulations, RNA molecules were modeled with the AMBER RNA force field^22^ by DES Research, while peptide fragments, water molecules, and ions were modeled with the amber99sb-disp force field^36^. Charges were neutralized with 0.1 M of NaCl. All production MDs were run in the NPT ensemble controlling temperature (T = 300 K) and pressure (P = 1 atm) respectively by means of the v-rescale^37^ and the Parrinello-Rahman^38^ schemes. Long-range electrostatic interactions were evaluated by using the particle mesh Ewald algorithm^39^ with a cutoff of 1 nm for the real space interactions, which is the most reliable and recommended scheme for dealing with long-range electrostatic interactions in highly-charged biomolecular systems^40^. Van der Waals interactions were computed with a cutoff distance of 1 nm. All bond lengths were constrained by using LINCS,^41^ allowing for the use of an integration time step of 2 fs. For each system three independent replicas were simulated for 1 μs starting from different initial configurations and velocities, saving solute conformations every 10 ps, while the first 200 ns were discarded from further analysis. Diffusion coefficients of peptide and oligonucleotide chains were estimated taking advantage of a recently published python tool by the Hummer group^42^.

### REST2 MD Simulations

REST2 simulations were run using GROMACS 2019.4^35^ patched with PLUMED 2.7,^43^ using a general Hamiltonian replica exchange implementation^.44^ 8 replicas were simulated, scaling the torsional, electrostatics, and Lennard-Jones potentials of the two solute molecules by factors in a geometric series between 1.0 and 0.7. A compensating charge was spread on the 6 Cl ions to maintain system neutrality. Replica exchanges were attempted every 400 steps. The acceptance rate was in the range 34%-39% for all pairs of neighboring replicas. Each replica was simulated for 480 ns. Only the unscaled replica was analyzed, discarding the first 80 ns.

### Generation of random configurations

Random configurations were generated with the following protocol: structures of R_5_, K_5_, and U_5_ molecules were extracted from MD simulations, discarding the first 200 ns, in order to form an ensemble of equilibrated configurations. An equal number of peptides (R_5_ or K_5_) and oligonucleotides (U_5_) were then randomly selected and placed in random orientations in a ∼145 nm^3^ rhombic dodecahedron box with the *gmx insert-molecules* tool, until the target concentration was reached (5, 10, and 20 molecules for each species at 125, 250, and 500 mg/ml, respectively).

### Analysis of the interactions

Two residues were considered in contact when at least one pair of heavy atoms was within a distance of 0.45 nm. In U_5_ molecules, O3’ and O5’ atoms were considered part of the phosphate group to which they were bound, thus the residue number of O3’ atoms were modified to correspond to those of the bound P atom. Surfaces of interaction for the simulations in highly-diluted conditions were computed with the *gmx sasa* tool in Gromacs 2018.3 as the difference between the sum of the surfaces of the two molecules and the total solvent accessible surface area (SASA) of the two molecules. Free energy profiles were computed with the standard equation F(S)=-k_B_T ln(P(S))+C, where P(S) is the probability distribution of the interaction surface between peptide and oligonucleotide molecules. The arbitrary constant C has been set to have the unbound state as reference (F(0)=0). The complex dissociation constants from REST simulations were estimated considering as bound states all conformations with at least one intermolecular contact between the two chains. Radial distribution functions (RDF) were estimated by means of the *gmx rdf* tool in Gromacs 2018.3. RDF between heavy atoms (Fig. 2D,E, and F) were computed considering all the intermolecular distances of non-hydrogen atoms of peptides and oligonucleotides. RDF between peptide side chains and RNA groups (Fig. 3C,D,E,F,G, and H), instead, were computed considering the distance between the terminal carbon atom of the side chains (CZ for arginine and CE for lysine, respectively) and the center of geometry of phosphate, sugar, and base groups. Hydrogen bond analysis was performed with the *gmx hbond* tool in Gromacs 2018.3 using the default definition of H-bond. The guanidinium group of arginine and the atoms of the nucleobases were considered to be forming a stacking interaction when the distance between the CZ atom of arginine and the center of geometry of the base ring is lower than 0.6 nm, and the angle between the plane of the guanidinium group and that of the base is lower than 30 degrees.

## Acknowledgments

We gratefully acknowledge the support by the French National Research Agency (ANR) under Grant ANR14-ACHN-0016 and the Swiss National Science Foundation under the grant CRSII5_193740.

## References

1. Shin Y, Brangwynne CP (2017) Liquid phase condensation in cell physiology and disease. Science 357:eaaf4382.

2. Banani SF, Lee HO, Hyman AA, Rosen MK (2017) Biomolecular condensates: organizers of cellular biochemistry. Nat. Rev. Mol. Cell Biol. 18:285–298.

3. Hyman AA, Weber CA, Jülicher F (2014) Liquid-Liquid Phase Separation in Biology. Annu. Rev. Cell Dev. Biol. 30:39–58.

4. Boeynaems S, Alberti S, Fawzi NL, Mittag T, Polymenidou M, Rousseau F, Schymkowitz J, Shorter J, Wolozin B, Van Den Bosch L, et al. (2018) Protein Phase Separation: A New Phase in Cell Biology. Trends Cell Biol. 28:420–435.

5. Weber SC, Brangwynne CP (2012) Getting RNA and Protein in Phase. Cell 149:1188– 1191.

6. Brangwynne CP, Tompa P, Pappu RV (2015) Polymer physics of intracellular phase transitions. Nat. Phys. 11:899–904.

7. Gomes E, Shorter J (2018) The molecular language of membraneless organelles. J. Biol. Chem.:jbc.TM118.001192.

8. Brady JP, Farber PJ, Sekhar A, Lin Y-H, Huang R, Bah A, Nott TJ, Chan HS, Baldwin AJ, Forman-Kay JD, et al. (2017) Structural and hydrodynamic properties of an intrinsically disordered region of a germ cell-specific protein on phase separation. Proc. Natl. Acad. Sci. 114:E8194–E8203.

9. Murthy AC, Dignon GL, Kan Y, Zerze GH, Parekh SH, Mittal J, Fawzi NL (2019) Molecular interactions underlying liquid-liquid phase separation of the FUS low-complexity domain. Nat. Struct. Mol. Biol. 26:637–648.

10. Wang J, Choi J-M, Holehouse AS, Lee HO, Zhang X, Jahnel M, Maharana S, Lemaitre R, Pozniakovsky A, Drechsel D, et al. (2018) A Molecular Grammar Governing the Driving Forces for Phase Separation of Prion-like RNA Binding Proteins. Cell 174:688-699.e16.

11. Garcia-Jove Navarro M, Kashida S, Chouaib R, Souquere S, Pierron G, Weil D, Gueroui Z (2019) RNA is a critical element for the sizing and the composition of phase -separated RNA–protein condensates. Nat. Commun. 10:3230.

12. Boeynaems S, Holehouse AS, Weinhardt V, Kovacs D, Lindt JV, Larabell C, Bosch LVD, Das R, Tompa PS, Pappu RV, et al. (2019) Spontaneous driving forces give rise to protein−RNA condensates with coexisting phases and complex material properties. Proc. Natl. Acad. Sci. 116:7889–7898.

13. Nott TJ, Petsalaki E, Farber P, Jervis D, Fussner E, Plochowietz A, Craggs TD, Bazett - Jones DP, Pawson T, Forman-Kay JD, et al. (2015) Phase Transition of a Disordered Nuage Protein Generates Environmentally Responsive Membraneless Organelles. Mol. Cell 57:936–947.

14. Gallego LD, Schneider M, Mittal C, Romanauska A, Gudino Carrillo RM, Schubert T, Pugh BF, Köhler A (2020) Phase separation directs ubiquitination of gene-body nucleosomes. Nature 579:592–597.

15. Ukmar-Godec T, Hutten S, Grieshop MP, Rezaei-Ghaleh N, Cima-Omori M-S, Biernat J, Mandelkow E, Söding J, Dormann D, Zweckstetter M (2019) Lysine/RNA-interactions drive and regulate biomolecular condensation. Nat. Commun. 10:2909.

16. Fisher RS, Elbaum-Garfinkle S (2020) Tunable multiphase dynamics of arginine and lysine liquid condensates. Nat. Commun. 11:4628.

17. Tsang B, Arsenault J, Vernon RM, Lin H, Sonenberg N, Wang L-Y, Bah A, Forman-Kay JD (2019) Phosphoregulated FMRP phase separation models activity-dependent translation through bidirectional control of mRNA granule formation. Proc. Natl. Acad. Sci. 116:4218– 4227.

18. Regy RM, Dignon GL, Zheng W, Kim YC, Mittal J (2020) Sequence dependent phase separation of protein-polynucleotide mixtures elucidated using molecular simulations. Nucleic Acids Res. 48:12593–12603.

19. Joseph JA, Espinosa JR, Sanchez-Burgos I, Garaizar A, Frenkel D, Collepardo-Guevara R (2021) Thermodynamics and kinetics of phase separation of protein-RNA mixtures by a minimal model. Biophys. J. 120:1219–1230.

20. Alshareedah I, Moosa MM, Raju M, Potoyan DA, Banerjee PR (2020) Phase transition of RNA−protein complexes into ordered hollow condensates. Proc. Natl. Acad. Sci. 117:15650–15658.

21. Paloni M, Bailly R, Ciandrini L, Barducci A (2020) Unraveling Molecular Interactions in Liquid–Liquid Phase Separation of Disordered Proteins by Atomistic Simulations. J. Phys. Chem. B 124:9009–9016.

22. Tan D, Piana S, Dirks RM, Shaw DE (2018) RNA force field with accuracy comparable to state-of-the-art protein force fields. Proc. Natl. Acad. Sci. U. S. A. 115:E1346–E1355.

23. Šponer J, Bussi G, Krepl M, Banáš P, Bottaro S, Cunha RA, Gil-Ley A, Pinamonti G, Poblete S, Jurečka P, et al. (2018) RNA Structural Dynamics As Captured by Molecular Simulations: A Comprehensive Overview. Chem. Rev. 118:4177–4338.

24. Wang L, Friesner RA, Berne BJ (2011) Replica exchange with solute scaling: A more efficient version of replica exchange with solute tempering (REST2). J. Phys. Chem. B 115:9431–9438.

25. Silmore KS, Howard MP, Panagiotopoulos AZ (2017) Vapour–liquid phase equilibrium and surface tension of fully flexible Lennard–Jones chains. Mol. Phys. 115:320–327.

26. Dignon GL, Zheng W, Kim YC, Best RB, Mittal J (2018) Sequence determinants of protein phase behavior from a coarse-grained model. PLOS Comput. Biol. 14:e1005941.

27. Ryan VH, Dignon GL, Zerze GH, Chabata CV, Silva R, Conicella AE, Amaya J, Burke KA, Mittal J, Fawzi NL (2018) Mechanistic View of hnRNPA2 Low-Complexity Domain Structure, Interactions, and Phase Separation Altered by Mutation and Arginine Methylation. Mol. Cell 69:465-479.e7.

28. Marsili S, Chelli R, Schettino V, Procacci P (2008) Thermodynamics of stacking interactions in proteins. Phys. Chem. Chem. Phys. 10:2673–2685.

29. Chavali SS, Cavender CE, Mathews DH, Wedekind JE (2020) Arginine Forks Are a Widespread Motif to Recognize Phosphate Backbones and Guanine Nucleobases in the RNA Major Groove. J. Am. Chem. Soc. 142:19835–19839.

30. Chong PA, Vernon RM, Forman-Kay JD (2018) RGG/RG Motif Regions in RNA Binding and Phase Separation. J. Mol. Biol. 430:4650–4665.

31. Wilson KA, Holland DJ, Wetmore SD (2016) Topology of RNA–protein nucleobase– amino acid π–π interactions and comparison to analogous DNA–protein π–π contacts. RNA 22:696–708.

32. Hajnic M, Osorio JI, Zagrovic B (2014) Computational analysis of amino acids and their sidechain analogs in crowded solutions of RNA nucleobases with implications for the mRNA–protein complementarity hypothesis. Nucleic Acids Res. 42:12984–12994.

33. Case DA, Cheatham TE, Darden T, Gohlke H, Luo R, Merz KM, Onufriev A, Simmerling C, Wang B, Woods RJ (2005) The Amber biomolecular simulation programs. J. Comput. Chem. 26:1668–1688.

34. Pettersen EF, Goddard TD, Huang CC, Couch GS, Greenblatt DM, Meng EC, Ferrin TE (2004) UCSF chimera - A visualization system for exploratory research and analysis. J. Comput. Chem. 25:1605–1612.

35. Abraham MJ, Murtola T, Schulz R, Páll S, Smith JC, Hess B, Lindahl E (2015) GROMACS: High performance molecular simulations through multi-level parallelism from laptops to supercomputers. SoftwareX 1–2:19–25.

36. Robustelli P, Piana S, Shaw DE (2018) Developing a molecular dynamics force field for both folded and disordered protein states. Proc. Natl. Acad. Sci. 115:E4758–E4766.

37. Bussi G, Donadio D, Parrinello M (2007) Canonical sampling through velocity rescaling. J. Chem. Phys. 126:014101.

38. Parrinello M, Rahman A (1981) Polymorphic transitions in single crystals: A new molecular dynamics method. J. Appl. Phys. 52:7182–7190.

39. Essmann U, Perera L, Berkowitz ML, Darden T, Lee H, Pedersen LG (1995) A smooth particle mesh Ewald method. J. Chem. Phys. 103:8577–8593.

40. York DM, Yang W, Lee H, Darden T, Pedersen LG (1995) Toward the Accurate Modeling of DNA: The Importance of Long-Range Electrostatics. J. Am. Chem. Soc. 117:5001–5002.

41. Hess B (2008) P-LINCS: A parallel linear constraint solver for molecular simulation. J. Chem. Theory Comput. 4:116–122.

42. Bullerjahn JT, von Bülow S, Hummer G (2020) Optimal estimates of self-diffusion coefficients from molecular dynamics simulations. J. Chem. Phys. 153:024116.

43. Tribello GA, Bonomi M, Branduardi D, Camilloni C, Bussi G (2014) PLUMED 2: New feathers for an old bird. Comput. Phys. Commun. 185:604–613.

44. Bussi G (2014) Hamiltonian replica exchange in GROMACS: a flexible implementation. Mol. Phys. 112:379–384.

