## supplemental material for "Arginine multivalency stabilizes protein/RNA condensates"

### Supporting Information

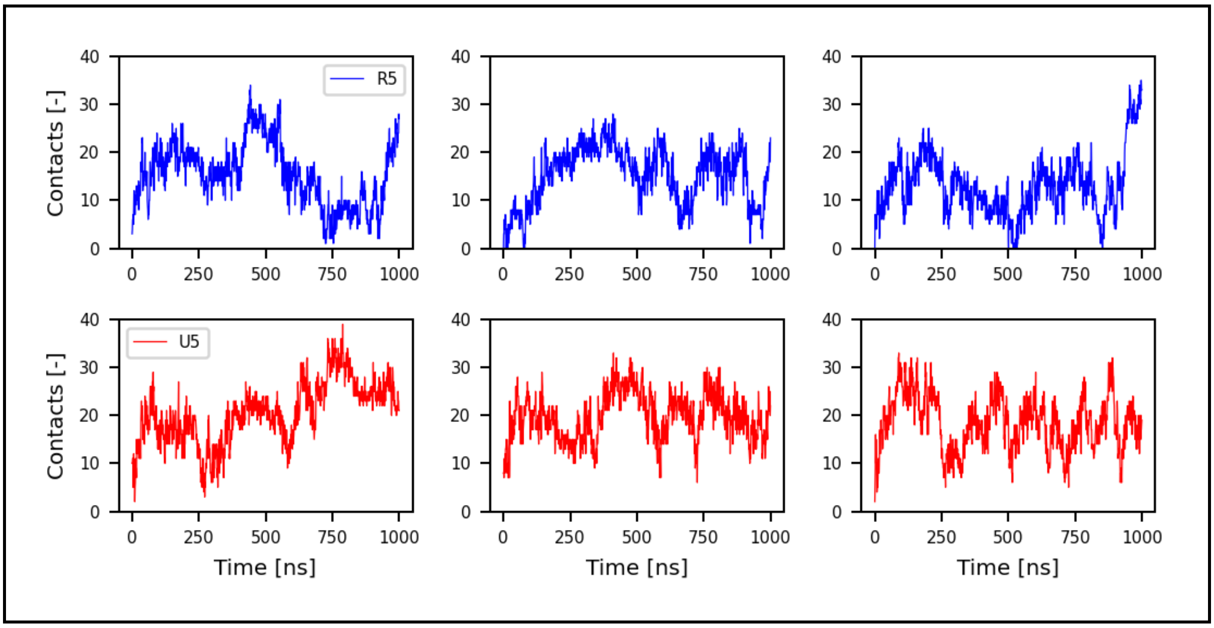


*Fig.S1 Representative timeseries of the number of intermolecular contacts of $R_5$ and $U_5$ molecules from MD simulations at high concentration.*

*Table S1. Average packing fraction (PF) of the high-density protein/RNA systems.*

| Protein/RNA Conc. [mg/ml] | <PF> [-] | σ [-] |
|---|---|---|
| $R_5/U_5$ | | |
| 125.0 | 0.127 | 0.003 |
| 250.0 | 0.222 | 0.004 |
| 500.0 | 0.342 | 0.031 |
| $K_5/U_5$ | | |
| 125.0 | 0.136 | 0.003 |
| 250.0 | 0.236 | 0.003 |
| 500.0 | 0.383 | 0.023 |
